## Supplemental Figures for "celldeath: a tool for detection of cell death in transmitted light microscopy images by deep learning-based visual recognition"

### Supplemental Figure 1

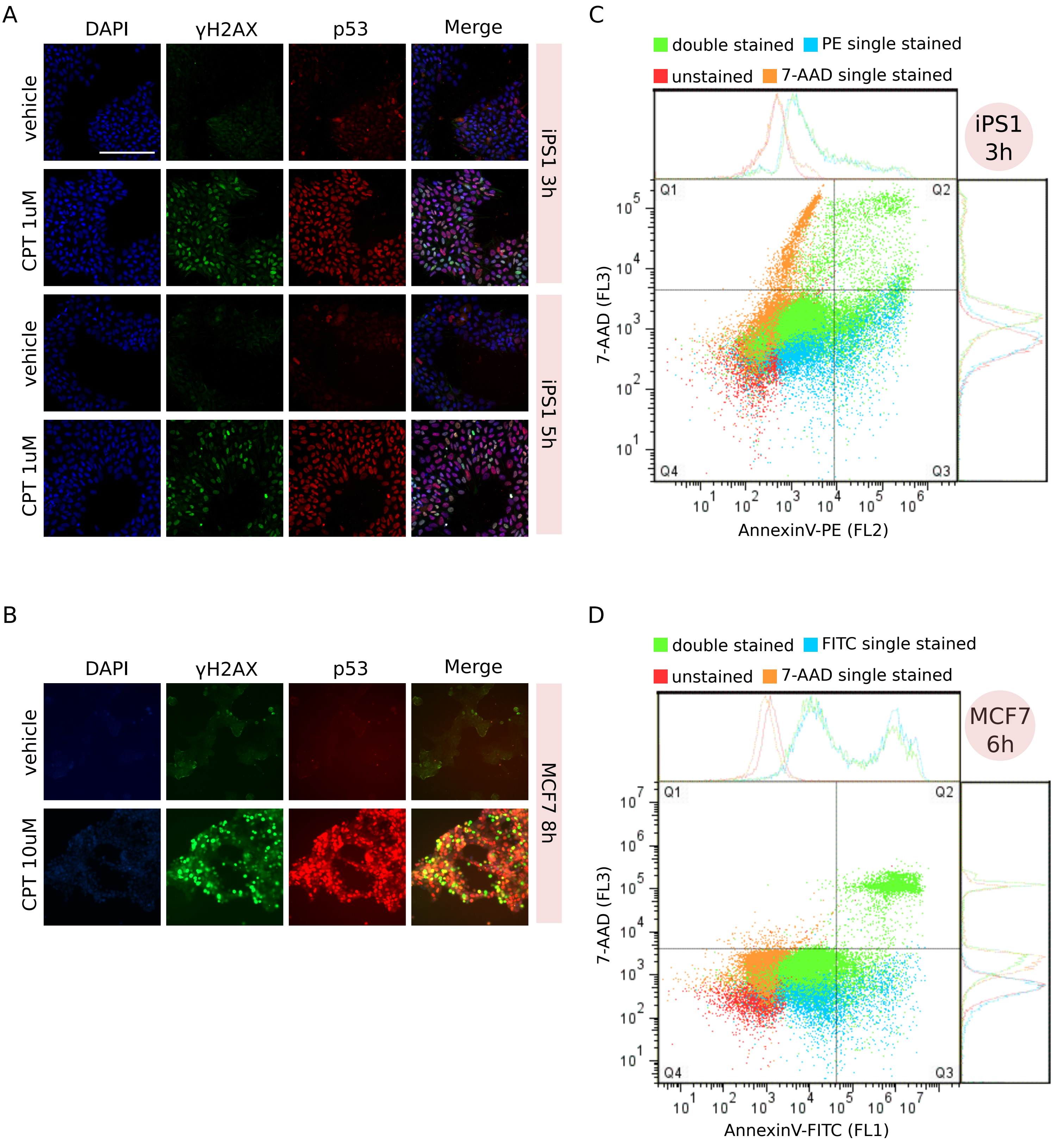

### Supplemental Figure 2

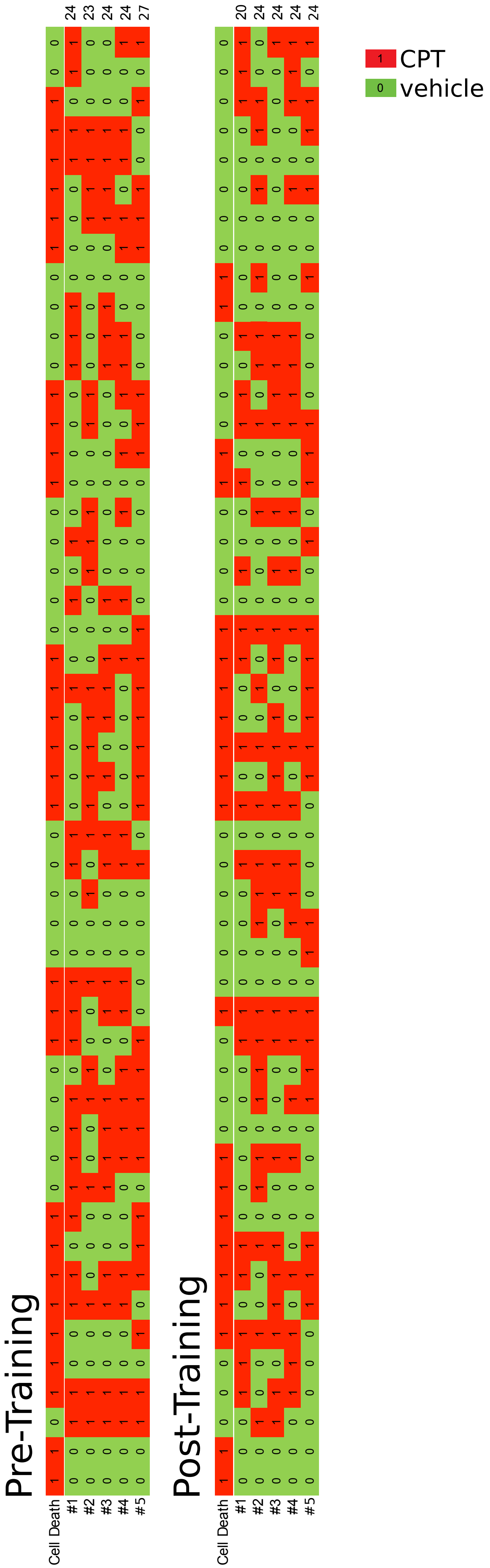

### Supplemental Figure 3

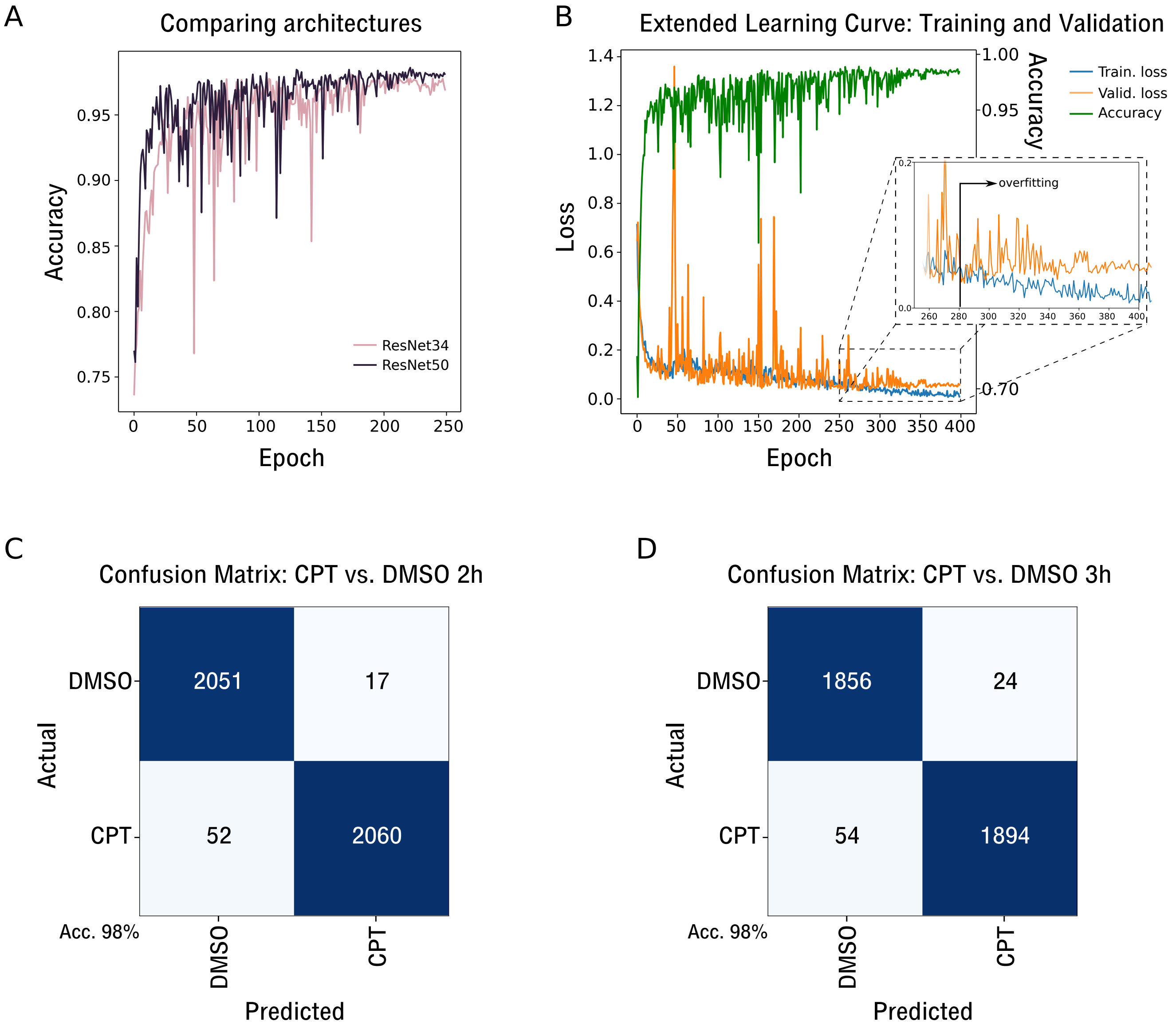

**S1 Table. Description of cell lines used in this work.**

| Name | Description | Species | Coating | Medium | Serum | Antibiotics | Origin |
|---|---|---|---|---|---|---|---|
| U2OS | Osteosarcoma cells | Human | No | DMEM | Yes | Yes | Dr. Martín Stortz |
| MCF7 | Luminal epithelial breast cancer cells | Human | No | DMEM | Yes | Yes | Dr. Luciano Vellón |
| T47D | Luminal epithelial breast cancer cells | Human | No | RPMI | Yes | Yes | Dr. Adalí Pecci |
| PC3 | Prostate cancer cells | Human | No | RPMI | Yes | Yes | Dr. Elba Vazquez |
| iPS1 | Induced pluripotent stem cell | Human | Geltrex | E8flex | No | No | Dr. Miriuka`s lab (Questa et al., 2016) |
| iPS2 | Induced pluripotent stem cell | Human | Geltrex | E8flex | No | No | Dr. Miriuka`s lab (Questa et al., 2016) |
| H9 | WA09 embryonic stem cell | Human | Geltrex | E8flex | No | No | Purchased from WiCell (USA) |

S2 Table. Deep learning model specifications

Sequential

| Layer (type) | Output Shape | Param # | Trainable |
|---|---|---|---|
| Conv2d | [64, 240, 320] | 9,408 | False |
| BatchNorm2d | [64, 240, 320] | 128 | True |
| ReLU | [64, 240, 320] | 0 | False |
| MaxPool2d | [64, 120, 160] | 0 | False |
| Conv2d | [64, 120, 160] | 4,096 | False |
| BatchNorm2d | [64, 120, 160] | 128 | True |
| Conv2d | [64, 120, 160] | 36,864 | False |
| BatchNorm2d | [64, 120, 160] | 128 | True |
| Conv2d | [256, 120, 160] | 16,384 | False |
| BatchNorm2d | [256, 120, 160] | 512 | True |
| ReLU | [256, 120, 160] | 0 | False |
| Conv2d | [256, 120, 160] | 16,384 | False |
| BatchNorm2d | [256, 120, 160] | 512 | True |
| Conv2d | [64, 120, 160] | 16,384 | False |
| BatchNorm2d | [64, 120, 160] | 128 | True |
| Conv2d | [64, 120, 160] | 36,864 | False |
| BatchNorm2d | [64, 120, 160] | 128 | True |
| Conv2d | [256, 120, 160] | 16,384 | False |
| BatchNorm2d | [256, 120, 160] | 512 | True |
| ReLU | [256, 120, 160] | 0 | False |
| Conv2d | [64, 120, 160] | 16,384 | False |
| BatchNorm2d | [64, 120, 160] | 128 | True |
| Conv2d | [64, 120, 160] | 36,864 | False |
| BatchNorm2d | [64, 120, 160] | 128 | True |
| Conv2d | [256, 120, 160] | 16,384 | False |
| BatchNorm2d | [256, 120, 160] | 512 | True |

| | | | |
|---|---|---|---|
| ReLU | [256, 120, 160] | 0 | False |
| Conv2d | [128, 120, 160] | 32,768 | False |
| BatchNorm2d | [128, 120, 160] | 256 | True |
| Conv2d | [128, 60, 80] | 147,456 | False |
| BatchNorm2d | [128, 60, 80] | 256 | True |
| Conv2d | [512, 60, 80] | 65,536 | False |
| BatchNorm2d | [512, 60, 80] | 1,024 | True |
| ReLU | [512, 60, 80] | 0 | False |
| Conv2d | [512, 60, 80] | 131,072 | False |
| BatchNorm2d | [512, 60, 80] | 1,024 | True |
| Conv2d | [128, 60, 80] | 65,536 | False |
| BatchNorm2d | [128, 60, 80] | 256 | True |
| Conv2d | [128, 60, 80] | 147,456 | False |
| BatchNorm2d | [128, 60, 80] | 256 | True |
| Conv2d | [512, 60, 80] | 65,536 | False |
| BatchNorm2d | [512, 60, 80] | 1,024 | True |
| ReLU | [512, 60, 80] | 0 | False |
| Conv2d | [128, 60, 80] | 65,536 | False |
| BatchNorm2d | [128, 60, 80] | 256 | True |
| Conv2d | [128, 60, 80] | 147,456 | False |
| BatchNorm2d | [128, 60, 80] | 256 | True |
| Conv2d | [512, 60, 80] | 65,536 | False |
| BatchNorm2d | [512, 60, 80] | 1,024 | True |
| ReLU | [512, 60, 80] | 0 | False |
| Conv2d | [128, 60, 80] | 65,536 | False |
| BatchNorm2d | [128, 60, 80] | 256 | True |
| Conv2d | [128, 60, 80] | 147,456 | False |
| BatchNorm2d | [128, 60, 80] | 256 | True |
| Conv2d | [512, 60, 80] | 65,536 | False |
| BatchNorm2d | [512, 60, 80] | 1,024 | True |

| | | | |
|---|---|---|---|
| ReLU | [512, 60, 80] | 0 | False |
| Conv2d | [256, 60, 80] | 131,072 | False |
| BatchNorm2d | [256, 60, 80] | 512 | True |
| Conv2d | [256, 30, 40] | 589,824 | False |
| BatchNorm2d | [256, 30, 40] | 512 | True |
| Conv2d | [1024, 30, 40] | 262,144 | False |
| BatchNorm2d | [1024, 30, 40] | 2,048 | True |
| ReLU | [1024, 30, 40] | 0 | False |
| Conv2d | [1024, 30, 40] | 524,288 | False |
| BatchNorm2d | [1024, 30, 40] | 2,048 | True |
| Conv2d | [256, 30, 40] | 262,144 | False |
| BatchNorm2d | [256, 30, 40] | 512 | True |
| Conv2d | [256, 30, 40] | 589,824 | False |
| BatchNorm2d | [256, 30, 40] | 512 | True |
| Conv2d | [1024, 30, 40] | 262,144 | False |
| BatchNorm2d | [1024, 30, 40] | 2,048 | True |
| ReLU | [1024, 30, 40] | 0 | False |
| Conv2d | [256, 30, 40] | 262,144 | False |
| BatchNorm2d | [256, 30, 40] | 512 | True |
| Conv2d | [256, 30, 40] | 589,824 | False |
| BatchNorm2d | [256, 30, 40] | 512 | True |
| Conv2d | [1024, 30, 40] | 262,144 | False |
| BatchNorm2d | [1024, 30, 40] | 2,048 | True |
| ReLU | [1024, 30, 40] | 0 | False |
| Conv2d | [256, 30, 40] | 262,144 | False |
| BatchNorm2d | [256, 30, 40] | 512 | True |
| Conv2d | [256, 30, 40] | 589,824 | False |
| BatchNorm2d | [256, 30, 40] | 512 | True |
| Conv2d | [1024, 30, 40] | 262,144 | False |

| | | | |
|---|---|---|---|
| BatchNorm2d | [1024, 30, 40] | 2,048 | True |
| ReLU | [1024, 30, 40] | 0 | False |
| Conv2d | [256, 30, 40] | 262,144 | False |
| BatchNorm2d | [256, 30, 40] | 512 | True |
| Conv2d | [256, 30, 40] | 589,824 | False |
| BatchNorm2d | [256, 30, 40] | 512 | True |
| Conv2d | [1024, 30, 40] | 262,144 | False |
| BatchNorm2d | [1024, 30, 40] | 2,048 | True |
| ReLU | [1024, 30, 40] | 0 | False |
| Conv2d | [256, 30, 40] | 262,144 | False |
| BatchNorm2d | [256, 30, 40] | 512 | True |
| Conv2d | [256, 30, 40] | 589,824 | False |
| BatchNorm2d | [256, 30, 40] | 512 | True |
| Conv2d | [1024, 30, 40] | 262,144 | False |
| BatchNorm2d | [1024, 30, 40] | 2,048 | True |
| ReLU | [1024, 30, 40] | 0 | False |
| Conv2d | [512, 30, 40] | 524,288 | False |
| BatchNorm2d | [512, 30, 40] | 1,024 | True |
| Conv2d | [512, 15, 20] | 2,359,296 | False |
| BatchNorm2d | [512, 15, 20] | 1,024 | True |
| Conv2d | [2048, 15, 20] | 1,048,576 | False |
| BatchNorm2d | [2048, 15, 20] | 4,096 | True |
| ReLU | [2048, 15, 20] | 0 | False |
| Conv2d | [2048, 15, 20] | 2,097,152 | False |
| BatchNorm2d | [2048, 15, 20] | 4,096 | True |
| Conv2d | [512, 15, 20] | 1,048,576 | False |
| BatchNorm2d | [512, 15, 20] | 1,024 | True |
| Conv2d | [512, 15, 20] | 2,359,296 | False |
| BatchNorm2d | [512, 15, 20] | 1,024 | True |
| Conv2d | [2048, 15, 20] | 1,048,576 | False |

| | | | |
|---|---|---|---|
| BatchNorm2d | [2048, 15, 20] | 4,096 | True |
| ReLU | [2048, 15, 20] | 0 | False |
| Conv2d | [512, 15, 20] | 1,048,576 | False |
| BatchNorm2d | [512, 15, 20] | 1,024 | True |
| Conv2d | [512, 15, 20] | 2,359,296 | False |
| BatchNorm2d | [512, 15, 20] | 1,024 | True |
| Conv2d | [2048, 15, 20] | 1,048,576 | False |
| BatchNorm2d | [2048, 15, 20] | 4,096 | True |
| ReLU | [2048, 15, 20] | 0 | False |
| AdaptiveAvgPool2d | [2048, 1, 1] | 0 | False |
| AdaptiveMaxPool2d | [2048, 1, 1] | 0 | False |
| Flatten | [4096] | 0 | False |
| BatchNorm1d | [4096] | 8,192 | True |
| Dropout | [4096] | 0 | False |
| Linear | [512] | 2,097,664 | True |
| ReLU | [512] | 0 | False |
| BatchNorm1d | [512] | 1,024 | True |
| Dropout | [512] | 0 | False |
| Linear | [2] | 1,026 | True |

Total params: 25,615,938
Total trainable params: 2,161,026
Total non-trainable params: 23,454,912
Optimized with 'torch.optim.adam.Adam', betas=(0.9, 0.99)
Using true weight decay as discussed in https://www.fast.ai/2018/07/02/adam-weight-decay/
Loss function : FlattenedLoss
======================================================================
Callbacks functions applied

**S3 Table. Number of images in each set for all conditions (related to Table 1 and Table 2).**

| Condition (1h) | Training and validation sets | Test set |
|---|---|---|
| CPT vs. DMSO | 11036 | 4188 |
| ALL vs. ALL | 11036 | 4188 |
| PC3 | 1472 | 508 |
| MCF7 | 1272 | 496 |
| T47D | 1316 | 480 |
| U2OS | 1424 | 828 |
| iPS1 | 2128 | 656 |
| iPS2 | 1492 | 496 |
| H9 | 1932 | 724 |
| PC3 out | 12972 | 2252 |
| MCF7 out | 13456 | 1768 |
| T47D out | 13428 | 1796 |
| U2OS out | 13244 | 1980 |
| iPS1 out | 12440 | 2784 |
| iPS2 out | 13236 | 1988 |
| H9 out | 12568 | 2656 |
